# The Mechanical Fingerprint of Hippocampal Sclerosis Linking Neuronal Cell Loss and Gliosis to Tissue Stiffness

**DOI:** 10.64898/2026.04.17.719271

**Authors:** Jan Hinrichsen, Nina Reiter, Lucas Hoffmann, Jörg Vorndran, Stefan Rampp, Daniel Delev, Oliver Schnell, Arnd Doerfler, Lars Bräuer, Friedrich Paulsen, Ingmar Blümcke, Silvia Budday

## Abstract

Hippocampal sclerosis (HS) is the most common pathology in drug-resistant temporal lobe epilepsy (TLE). However, clinical diagnosis, prevalent epileptogenicity, and drug drug-resistance in individuals with HS remain an ongoing challenge demanding multidisciplinary research efforts. In this study, we examined the mechanical properties of neurosurgically en bloc resected HS specimens (n=8) *ex vivo* under compression, tension, and torsional shear. We fitted a two-term Ogden hyperelastic model to the measured mechanical responses to quantify nonlinear mechanical tissue properties. The resulting parameters revealed higher strain stiffening under compression in HS compared to hippocampus obtained post mortem (n=7). The distinction was most noticeable in the large-strain regime, which has important implications for using mechanical tissue properties as valuable diagnostic biomarker. Furthermore, we correlated the tissue microstructure with mechanical parameters. We trained a deep-learning histopathology classifier to detect and classify neurons and glial cells from hematoxylin-stained whole slide images (WSI). We identified a strong association between the small-strain stiffness (shear modulus *µ*) and the overall cell density as well as the glial cell density. The negative relationship between the neuron-to-glia ratio and shear modulus is consistent with the hypothesis that neuronal cell loss and gliosis drives tissue stiffening, respectively. Magnetic resonance imaging (MRI) analysis of the specimens confirmed the previously reported negative association between MRI-derived fractional anisotropy and shear modulus *µ*. Taken together, our study establishes a direct link between tissue mechanics and microstructure, suggesting nonlinear continuum mechanics models as promising new tools for clinical diagnosis and novel research strategies.

## Introduction

HS is the most prevalent structural brain lesion in individuals suffering from drug-resistant focal epilepsy amenable to selective neurosurgical resection [6], with approx. 70% of individuals being seizure free five years after surgery [44]. Hippocampal sclerosis (HS) is histopathologically defined by the International League against Epilepsy (ILAE) as segmental neuronal cell loss in the Ammon’s horn with concomitant reactive gliosis [7]. These alterations can be presurgically recognized in approx. 90% of individuals by using dedicated magnetic resonance imaging (MRI) protocols, i.e., HARNESS [5], demonstrating atrophic volume loss, and increased T2-FLAIR signaling as characteristic hallmarks of HS [53, 75]. Brain somatic mutations in genes associated with the MAP-Kinase signaling pathway, e.g., PTPN11 [39], and/or human herpes virus infections, e.g., HHV6 [1], are likely pathogenic causes of HS.

Part of the pre-surgical planning is the proper hypothesis on the location of the epileptogenic zone that needs to be removed in order to achieve seizure freedom. The planned resection needs to respect the epileptological-functional balance aiming at the tissue causing seizure without causing neurological deficit. While electroencephalography (EEG) and magnetoencephalography (MEG) measurements are valuable diagnostic methods for the detection and location of the epileptogenic focus, they are often limited in the achievable accuracy [3]. Invasive methods like intracranial electroencephalography (iEEG) enable higher achievable sensitivity using electrodes that are implanted into the patient’s brain [61]. Still, this accuracy comes with a significant burden of the highly invasive method, which motivates the development of alternative diagnostic approaches.

The term sclerosis stems from the ancient greek 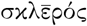 (sklē rós) ‘hard’ and a sclerotic hippocampus will appear atrophied and harder in intraoperative palpation [58]. Quantifying these mechanical properties promises to enable new diagnostic methods, e.g. based on force sensing surgery instruments, to detect and locate HS. In recent decades, research focusing on the biomechanical properties of brain tissue has gained significant momentum [73, 14]. Brain mechanics is an important research domain to understand the biophysical response of a given anatomical brain region, e.g., during development [26, 70], crash scenarios [22, 48] and surgical intervention [38, 54] amongst many other use cases. Also at the cell scale, mechanics plays an important role, e.g in axonal path finding [42], mechano-sensitive ion channels [73, 42], and regulation of neurotransmission and synaptic activity [74, 77].

From a mechanical perspective, brain tissue behaves highly nonlinear under large deformations. Especially characteristic is the higher strain stiffening in compression than in tension, denoted as compression-tension asymmetry [12]. These properties necessitate material models based on nonlinear continuum mechanics that are able to capture such complex effects. During a neurosurgical procedure, large brain tissue deformations are common and these complex effects significantly influence how the tissue reacts to mechanical inputs and how it feels to the surgeons hands.

Most studies have focused on characterizing the mechanical behavior without pathological conditions [23] and continued efforts to understand and accurately model mechanical effects like conditioning behavior [13], region-specific properties [33], as well as the linking to tissue constituents [46, 32] and *in vivo* imaging data [72] promise to enable highly accurate and spatially resolved simulation models. By additionally considering pathological conditions, mechanical modeling and characterization could enable the identification of mechanical thresholds that distinguish pathological from healthy conditions, as a promising new diagnostic tool to locate pathologies. Previous studies on the mechanical characterization of pathological human brain tissue have largely focused on tumors [19, 68, 18, 60, 20, 56]. In animals, researchers have investigated the mechanical changes in rat brain tissue due to stroke [50] or acidosis [34]. More recently, there have also been studies aiming to determine mechanical properties of epileptic human brain tissue using *ex vivo* indentation [25, 59].

However, the distinct changes in mechanics as they are felt by surgeons in sclerotic hippocampal tissue have not yet been investigated through biomechanical testing and analyses. In this study, we measure the *ex vivo* mechanical response of surgically resected HS tissue as well as controls from body donors. A two-term Ogden model enables us to quantify the nonlinear mechanical properties and identify the “mechanical fingerprint” of HS marked by distinct compression stiffening. We introduce a neural network cell classifier that enables us to automatically process stained whole slide images of the tested tissue and obtain its composition in terms of cellular densities. By combining cellular densities and mechanical properties, we are able to relate changes in tissue stiffness to neuronal cell loss and gliosis.

## Methods

### Human brains from body donors

As non-HS controls we used hippocampus samples from post mortem human brains. We obtained 11 brains (4 female, 7 male) from body donations and one brain (male) from autopsy. The age of the donors ranged from 57 to 92 years (mean age 71.1 years). After extraction from the skull, we first measured the brains in an MRI scanner and then transported them in buckets filled with artificial cerebrospinal fluid (ACSF, for constituents see [33]) to our biomechanics laboratory for mechanical testing.

For the donor brains, exact post mortem times are known. We received the donors less than 24 hours post mortem. The post mortem intervals at the time of the mechanical experiments range from 26 to 89 h (mean 55.5 h) for the individual samples. For the autopsy brain, only the day but not the exact time of death is known. The post mortem interval at the time of the autopsy was maximum 48 hours and the mechanical experiments for hippocampus samples of this brain were performed shortly after autopsy.

Before starting mechanical experiments, we sectioned the brains in the coronal direction. Coronal slices were kept refrigerated in ACSF-filled containers until the respective slice was used for mechanical testing. We used a circular punch to extract samples of `8 mm` diameter.

### Surgical hippocampus samples

We used surgically resected tissue from 7 patients that underwent epilepsy surgery. The age of the patients ranged from 21 to 55 years (mean age 35.7 years) and all of them were diagnosed with hippocampus sclerosis type I after histopathological analysis. A part of the resected tissue that was not needed for diagnostics was reserved for mechanical testing, placed in an ACSF-filled tube, and transported to the biomechanics laboratory immediately after resection. The slice for mechanical testing usually already had the required thickness of approximately `4 mm`, so a cylindrical sample could directly be punched from the received tissue slice. Whenever possible, we tried to obtain samples of `8 mm` diameter. However, we used a `6 mm` punch for one sample due to its small size.

### Obtaining fractional anisotropy and hippocampi volume ratios from MRI data

Body donor brains were scanned embedded in ACSF within a 3D-printed container device [40]. Diffusion-weighted images (DWI) were acquired on a 3 T Magnetom Prisma scanner and a 3 T Magnetom Cima.X scanner (Siemens Healthineers, Forchheim, Germany) using a 2D EPI diffusion sequence with parameters depending on the scanner: Prisma: TE= 69 ms, and TR= 4100 ms. The slice thickness was 4 mm and spacing between slices was 5.2 mm. The in-plane resolution was 1.71875 mm x 1.71875 mm. The flip angle was 90^◦^. The pixel bandwidth was 1955 Hz. A DTI scheme was used, and a total of 20 diffusion sampling directions were acquired three times for improved SNR at a b-value of 1000 s*/*mm^2^. Cima.X: TE= 44 ms, and TR= 3500 ms. The slice thickness was 2 mm. The spacing between slices is 2.6 mm. The in-plane resolution was 1.71875 mm x 1.71875 mm. The flip angle was 90^◦^. The pixel bandwidth was 1698 Hz. A DTI diffusion scheme was used, and a total of 30 diffusion sampling directions were acquired twice. The b-value was 1000 s*/*mm^2^. The tensor metrics were calculated using DWI with b-value lower than 1750 s*/*mm^2^. In addition, a 3D T1-weighted (T1w) MPRAGE was acquired with an isotropic voxel resolution of 1 mm (TI= 900 ms, flip angle 9◦). Fractional anisotropy (FA) was then calculated based on the DWI data with manufacturer’s software. T1w images were processed with the Computational Anatomy Toolbox (CAT12) [24] included in brainstorm [69], which is documented and freely available for download online under the GNU general public license^1^. The segmentation procedure fits several anatomical atlases and parcellation schemes to the individual MRI, of which the Hammers atlas [28] was utilized to extract the hippocampal volume, volume ratios, and mean FA over all voxels contained within the hippocampi.

### Mechanical testing

For mechanical characterization of the tissue samples, we used DHR-3 and HR-30 rheometers from TA Instruments (New Castle, Delaware, USA). For sensitivity specifications of the rheometers, the reader is referred to [66]. We subjected the samples to cyclic loading and stress relaxation experiments in compression, tension, and shear as shown in Table 1 and illustrated in Figure 1 h).

**Table 1.** Testing protocol.

|  |  |  |
| --- | --- | --- |
| 1 | **Frequency sweep** | at 1% strain |
| 2 | **Cyclic compression/tension in z-direction** | up to 15% strain |
| 3 | **Compression relaxation in z-direction** | at 15% strain; hold time=300s |
| 4 | **Tension relaxation in z-direction** | at 15% strain; hold time=300s |
| 5 | **Cyclic torsional shear** | 3 Cycles up to a shear of $\gamma = 0.15$ |
| 6 | **Cyclic torsional shear** | 3 Cycles up to a shear of $\gamma = 0.3$ |

**Figure 1.**
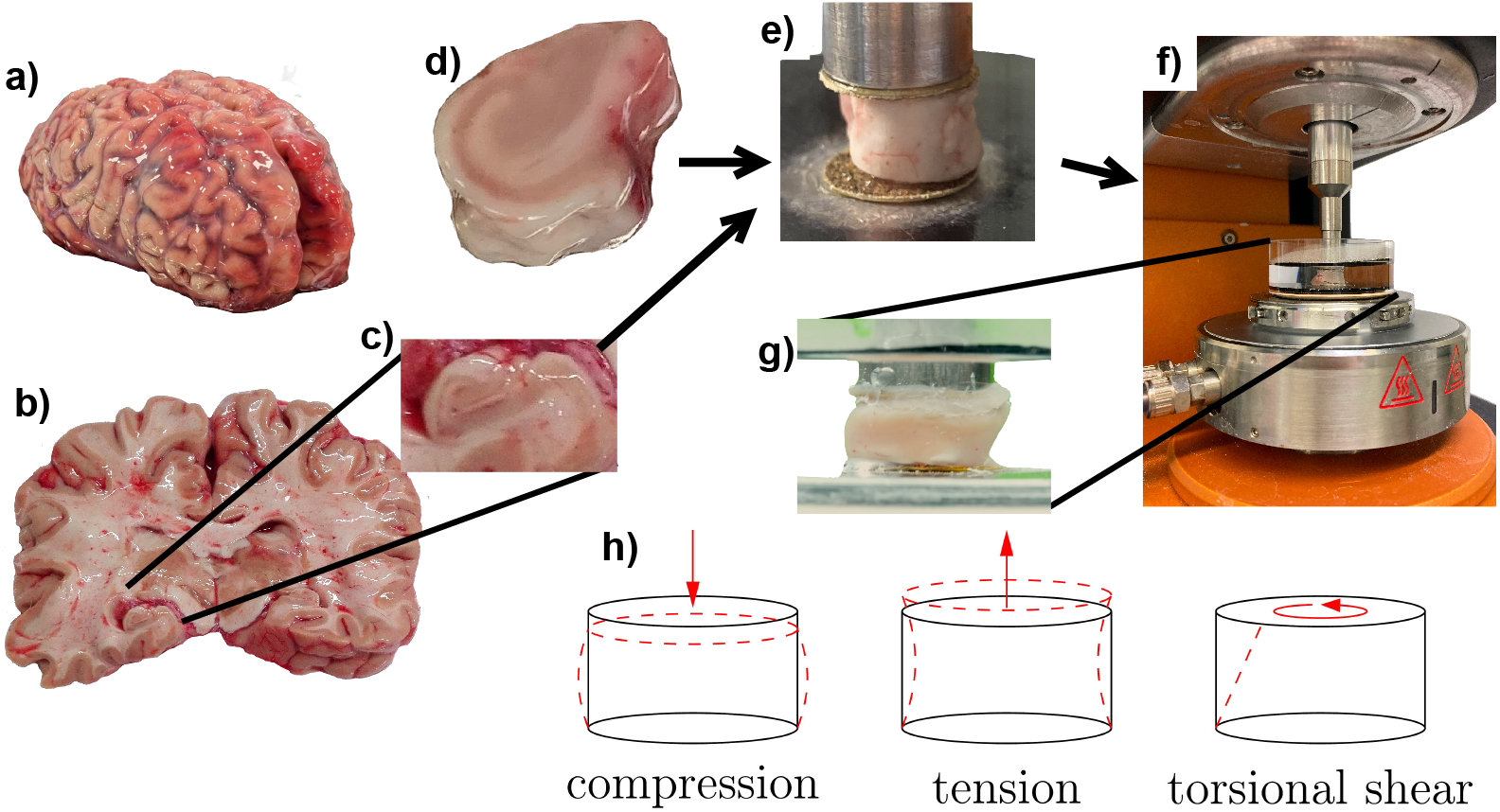
Specimen extraction and experimental setup. a,b,c) Hippocampus from body donor. d) Surgically resected hippocampus tissue. e,f) Sandpaper is glued to the rheometer specimen holders and then the extracted specimens are glued to the sandpaper. During testing, specimens are submerged in phosphate buffered solution (PBS) or artificial cerebrospinal fluid (ACSF). g) Submerged specimen during testing. h) Loading modes tested in the rheometer.

To prepare the rheometer, we glued sandpaper to the upper and lower specimen holders and calibrated the rheometer with the sandpaper attached. Subsequently, we glued the sample to the upper and lower sample holder using super glue. We attached the sample first to the upper sample holder and then carefully lowered it until it was in full contact with the glue on the lower sample holder but not yet pre-compressed. The testing setup is shown in Figure 1 e to g). Before starting the experiment, we immersed the sample in phosphate-buffered saline solution and heated the Peltier plate (below the lower sample holder) to 37 ◦C to mimic physiological conditions. After the experiment, we carefully extracted the sample with a scalpel and fixed it in 4% formalin for subsequent histological analysis.

We obtained the raw data recorded by the rheometer using the data logging application (ARG2AuxiliarySample.exe) provided by TA Instruments and located in the TA Instruments Trios installation folder. We used force, gap, torque and encoder data to calculate the stresses and strains in the sample. For compression and tension tests, the stretch is calculated as *λ* = *h/H* = [*H* + Δ*z*]*/H*, where *H* is the original height of the sample and *h* is the current gap height, i.e., *h < H* in compression tests and *h > H* in tension tests. The nominal stress *P* is obtained by dividing the measured force *F* through the sample’s undeformed cross section area: *P* = *F/A*. For torsional shear tests, we obtain the current shear strain from encoder data (angle Θ in rad) as *γ* = *R \** Θ*/H* and the shear stress as *τ* = 2*t/πR*^3^, where *t* is the measured torque and *R* is the radius of the sample.

Torque data contained unphysical spikes at the beginning of the curves that are artifacts stemming from the torque sensor’s inertia. These spikes have been excluded by z-score filtering as well as thresholding in comparison to moving average and Savitzky-Golay filter results. The data have been processed using a moving average filter. Finally, a Ramer–Douglas–Peucker algorithm has been applied to reduce the number of data points without losing too much information.

Figure 2 a,b shows the measured mechanical response of one specimen. After data processing, the data shown in Figure 2 c,d is obtained, along with the approximated hyperelastic response, obtained by averaging the loading and unloading data.

**Figure 2.**
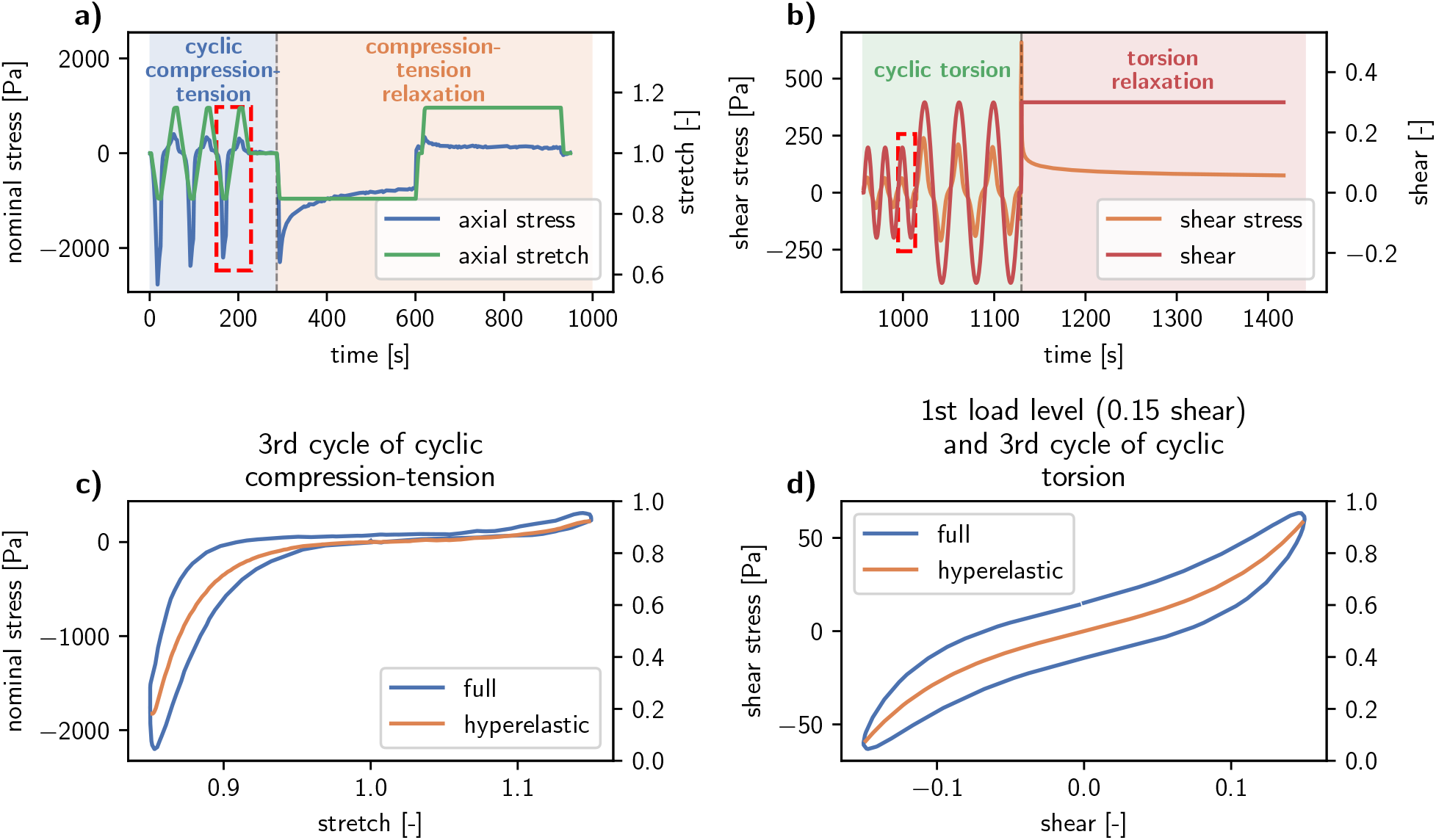
Measured mechanical data and hyperelastic response extraction: a-b) show the measured mechanical response for one specimen for the whole protocol in Table 1. The data is split into two consecutive parts starting with a) compression-tension loading modes with cyclic and relaxation loading where nominal stress and stretch in axial direction are shown and followed by b) torsional shear loading modes with cyclic and relaxation loading where torsional shear stress and the torsional shear are shown. Red boxes in a) and b) mark the loading cycles that were used for mechanical characterization and are shown in c) and d), respectively. c) Third cycle of the cyclic compression-tension loading mode with the viscoelastic hysteretic (full) as well as the approximated hyperelastic response, obtained by averaging loading and unloading. d) The same hyperelastic approximation for the third cycle of the first loading mode of cyclic torsional loading.

### Mechanical modeling

To characterize the mechanical response, we use a parameterized mechanical model to approximate the measured force-displacement data. Then, for each specimen, we identify the set of parameters that minimizes the error between the mechanical model output and the measured experimental data. This enables us to quantitatively compare specimens. For the material model describing the relationship between stress and strain, we use a constrained, two-term, hyperelastic Ogden model. By selecting a hyperelastic model, we assume that the mechanical response exhibits no history dependency and follows the same stress-strain path during loading and unloading. Although brain tissue exhibits viscoelastic behavior, these effects depend on the loading rate and can be neglected for slow deformation changes, such as those occurring during palpation or surgical interventions targeted by our study. Ogden-type models have been shown to capture the mechanical behavior of brain tissue, especially the pronounced compression-tension asymmetry characteristic of higher strain stiffening in compression than in tension [12].

The deformation map *φ* : **X** → **x** maps points from the undeformed material configuration **X** to the deformed configuration **x**. From the deformation gradient **F**, defined as Grad *φ*(**X**), we obtain the principal stretches *λ*_*a*_ as square root of the principal values of the left Cauchy-Green tensor **b** = **FF**^T^ [57]. The constitutive relation is given in terms of the strain energy-density Ψ. For hyperelastic materials, Ψ depends only on the deformation gradient **F** or its derived strain tensors. Under the assumption of incompressibility we can then calculate the first Piola-Kirchhoff stress

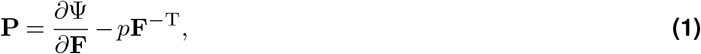

where p acts as Lagrange multiplier and is also denoted as pressure [35].

We choose a two-term Ogden model and reformulate it in terms of the shear modulus *µ*. The generalized Ogden model with N terms is defined as

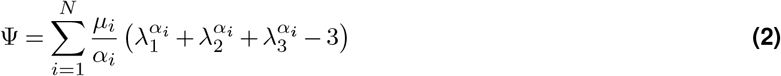

with the classical shear modulus given by 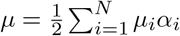 [35]. For our two-term model, we now constrain the exponents *α*_1_ ≤ 0 and *α*_2_ ≥ 0. This will cause the first term to capture the compression tension asymmetry with higher strain stiffening in compression, controlled by *α*_1_. The second term then captures the strain-stiffening effects in tension controlled by *α*_2_ . By defining 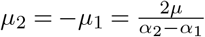 we can then reformulate Equation 2 to obtain

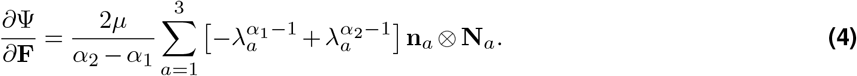

Subsequently, 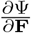 in Equation 1 can be rewritten using the spectral decomposition **F** = *λ*_*a*_**n**_*a*_ ⊗ **N**_*a*_ as

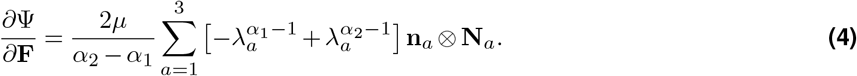

The rheometer used as mechanical testing device prescribes the deformation of our approximately cylindrical specimens in axial directions and measures the mechanical response in terms of the axial force acting on the specimen holder of the rheometer. These data is not enough to fully describe the deformation and stress state of our specimens. Therefore, we need to include additional assumptions into our mechanical model to obtain a fully defined deformation and stress state. This enables us to determine the parameters for the chosen constitutive equation Equation 3. We assume the specimens to have perfect cylindrical geometries with the diameter of the punching tools and the height determined from the measured data. During loading we further assume the specimens to deform homogeneously in compression and tension, while the shear *γ* for simple torsional shear depends on the radius *γ* = *γ*(*r*). The mechanical response is assumed to be hyperelastic, isotropic, and homogeneous throughout the specimen geometry. Uniaxial compression tension loading is then described by the deformation gradient

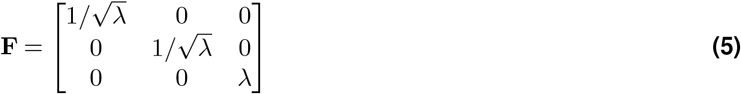

with the principal stretches 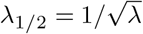 and *λ*_3_ = *λ*. The axial stress *P*_1_ is then obtained from Equation 1 and Equation 4, where p is calculated from the condition *P*_*x/y*_ = 0. Considering the assumptions made before, this is equal to 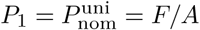, which we denote as nominal stress, with the axial force *F* and the undeformed area *A* of the ideal cylindrical geometry. The shear *γ* in a cylinder with height H, that is loaded with the torsion angle *φ*, is obtained as *γ*(*r*) = *φ * r/H*. Thus, we obtain the deformation gradient F in cylindrical coordinates, following the derivations in [30], as

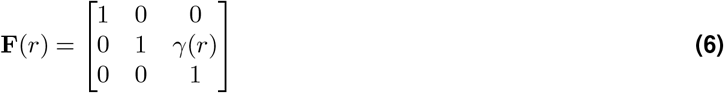

with the principal stretches *λ*_1_ = 1 and 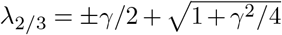. Subsequently, we use a numerical Gauss integration scheme with four integration points and obtain the torque T by integrating the shear stress

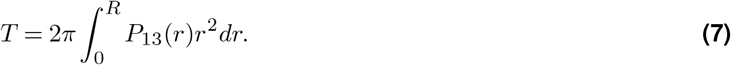

Then we can calculate the nominal shear stress in terms of the maximum stress at *r* = *R* as 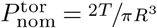. Figure 3 shows the influence of the three model parameters *µ, α*_1_ and *α*_2_ on the predicted mechanical response in compression-tension and torsional shear. *µ* controls the linear part of the model, shown in Figure 3 a,d, and especially the response for small deformations. The nonlinearity is controlled by the *α* parameters with *α*_1_ controlling the compression stiffening behavior in Figure 3 b,e and *α*_2_ the tension stiffening behavior in Figure 3 c,f. Higher absolute values for *α*_2_ are needed to cause the same (absolute) change of maximum predicted stresses in tension as for *α*_1_ in compression.

**Figure 3.**
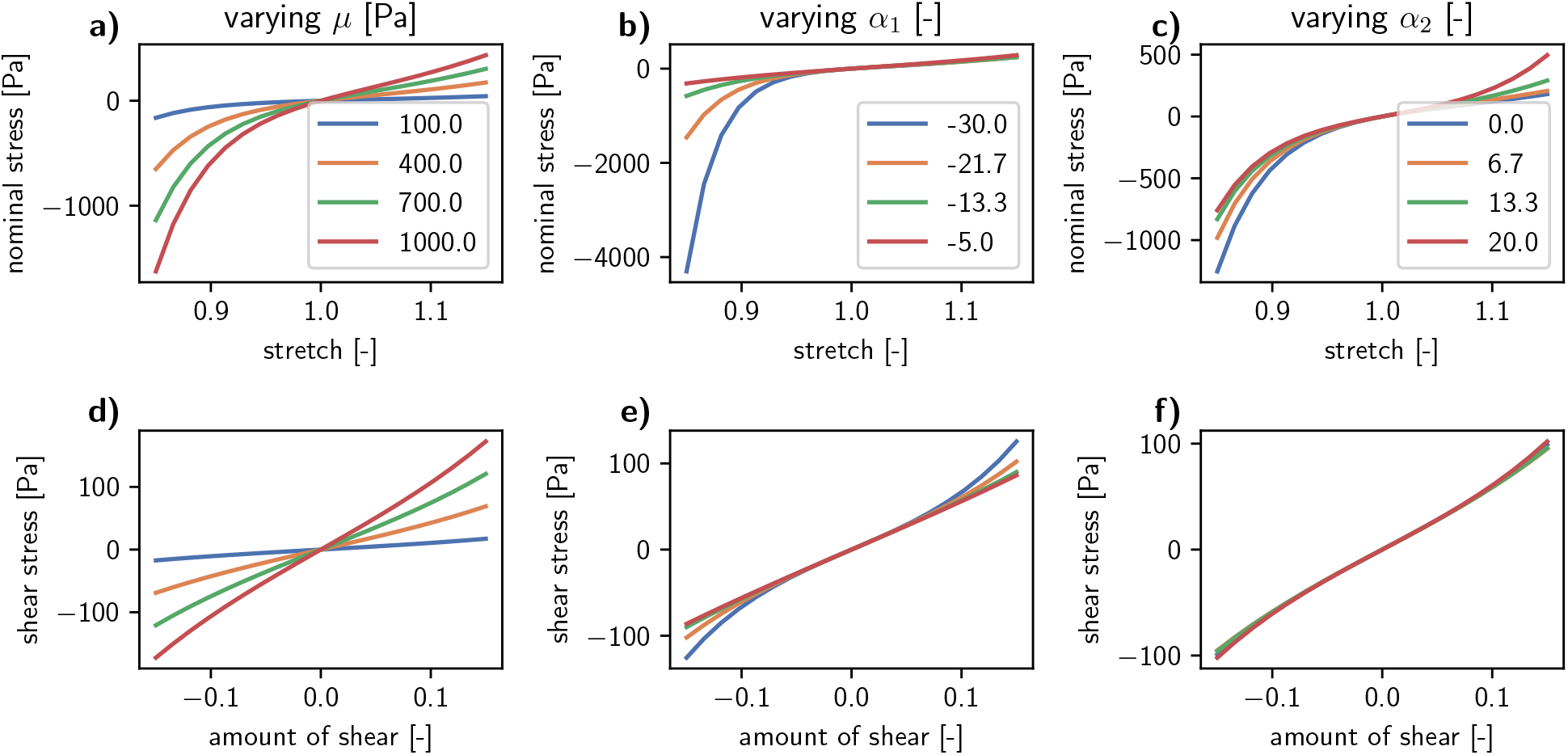
Parameter study showing the influence of the three model parameters *µ, α*_1_, and *α*_2_ on the predicted mechanical response: a-c) Predicted response under compression-tension loading in terms of the predicted nominal stresses for prescribed stretches. d-f) Predicted response under torsional shear loading in terms of predicted shear stress under prescribed shear. a,d) Influence of the shear modulus *µ*. b,e) Influence of the compression stiffening *α*_1_. c,f) Influence of the tension stiffening *α*_2_

The parameter identification scheme uses a trust region reflective algorithm to minimize the difference between experimental measurement data and predicted model output. All loading modes—compression, tension, and torsional shear—are fitted simultaneously to obtain the generalized material response. Further details on the parameter identification scheme can be found in our previous study [33].

### Deep neuropathology - Cell classifier

#### Data acquisition and preprocessing

The training dataset consisted of 28 hematoxylin-stained, digitized hippocampal tissue sections obtained from 16 individuals, encompassing both samples with neuropathologically confirmed hippocampal sclerosis and non-sclerotic hippocampal controls. Neocortical tissue samples provided additional control data. The cohort included both surgically resected tissue specimens and post-mortem samples. To ensure precise identification of cellular structures within these heterogeneous samples, we systematically classified cells into eleven distinct classes and two technical auxiliary categories listed in Appendix 3. Annotations were performed by an experienced expert (IB) using the NDPI-Viewer software (Hamamatsu Photonics Europe) to define bounding boxes for each cell type. Finally, RGB image patches (200 µm edge length, 907 x 907 pixel resolution) were generated using a random-shifting procedure to ensure robust sampling for the subsequent model.

#### Segmentation and mask generation

To refine the annotated image segments, we employed a combination of adaptive filtering, thresholding, and morphological operations, integrated with an object recognition workflow based on a local instance of the Segment Anything Model (SAM)[41]. Using a custom Python pipeline, we generated training masks corresponding to distinct cell types. While each mask targeted an explicitly labeled object, the segments inherently included adjacent cellular structures—comprising both verified annotations and algorithmically predicted instances. By preserving these overlapping structures and complex backgrounds, we sought to capture the natural heterogeneity of the samples, thereby enhancing the robustness and generalization of the detection model.

#### Dataset statistics and augmentation

We aimed for a minimum of 2,000 annotations per cell class to ensure statistical depth. While most classes met this threshold, the CAY (corpora amylacea, n = 1,259) and EDY (ependymal cell, n = 786) cohorts remained below the target. The GC (granule cell) class exhibited the highest annotation density (n = 5,370), whereas EDY (ependymal cell) represented the least frequent class in the raw dataset. To mitigate class imbalance and enhance model stability, we expanded the dataset by 20,000 mask-image pairs using a pseudo-labeling and copy-and-paste augmentation strategy. This involved systematically extracting validated annotations and cell segments and integrating them into novel background contexts derived from the original native imagery. This targeted synthesis significantly improved the representation of minority classes while preserving the biological authenticity of the image backgrounds.

#### Computational model

We implemented a Mask R-CNN architecture using the PyTorch framework [62] for automated cell detection and instance segmentation. The model utilized a ResNet-50 backbone [31], pre-trained on the COCO dataset [45], to facilitate robust feature extraction. The objective function integrates the errors from the Region Proposal Network (RPN)—specifically the RPN class and bounding box losses—alongside the final stage losses for classification, localization, and mask segmentation (LRPN + LOBJ + LCLS + Lbox + Lmask). To enhance performance, the objective function employed a modified cross-entropy loss, specifically weighted to account for the observed class distribution and dataset structure. The training process utilized a dynamic learning rate of 0.0005 to ensure stable convergence. To further enhance classification accuracy and filter out auxiliary technical artifacts, we integrated a post-processing patch into the workflow.

Training was strictly regulated via an early-stopping criterion based on the validation loss to prevent overfitting and ensure robust generalization. The final model reached convergence after 14 epochs and the final multi-validation loss settled at 0.322.

We obtained cellular densities by calculating the tissue area in each whole slide image (WSI) using the Otsu’s thresholding method provided by the OpenCV library [9].

#### Validation

Figure 4 shows the results of the cell detection classifier on sections of hematoxylin-stained hippocampus specimens. The classifier can detect and classify most visible nuclei.

**Figure 4.**
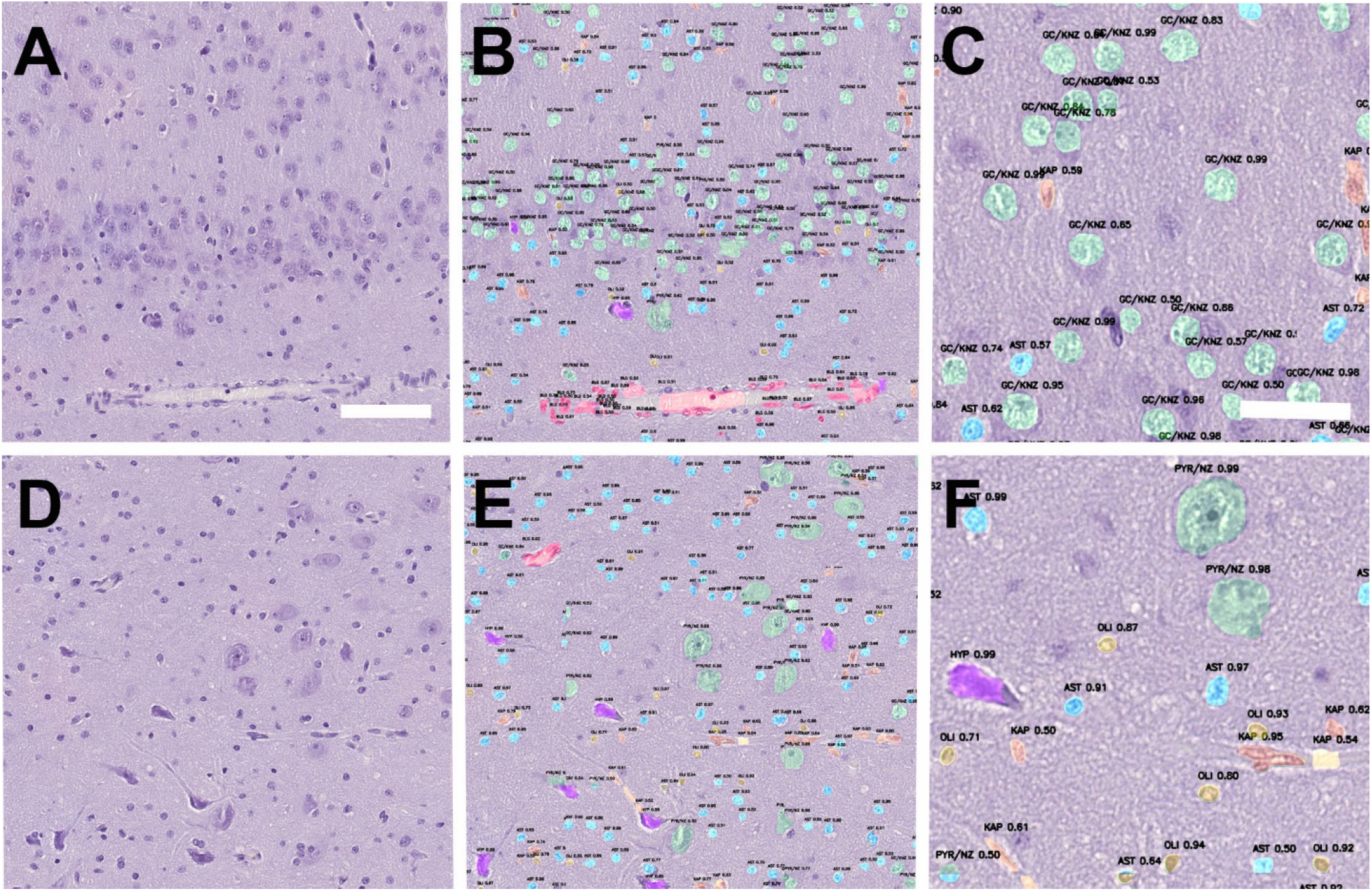
A deep-learning based cell segmentation classifier developed in this study: hematoxylin-stained section of the human dentate gyrus (surgical specimen with granule cell dispersion). B: same region from A showing the automated cell annotation of the novel cell segmentation classifier. C: Higher magnification of region shown in A and B, illustrating the granularity of information provided by the classifier. D: hematoxylin-stained section of the CA4 sector of the human hippocampus (surgical specimen). E: same region from D showing the automated cell annotation. F: Higher magnification of region shown in D and E. PYR/GC = pyramidal and granule cells (green); HYP = hypoxic neurons (violet); AST = astrocytes (blue); OLI = oligodendrocytes (yellow); KAP = endothelial cells of capillaries (orange). Numbers represent the confidence score for each annotated cell class. Scale bar in A = 100 µm, applies also to B, D, E. Scale bar in C = 50 µm, applies also to F.

A custom validation framework provides a quantitative evaluation of the predictive performance, overcoming the limitations of Intersection over Union due to incomplete mask labeling. This approach quantifies model performance by measuring recall against manual annotations, as well as using a discovery metric to account for unlabeled cells identified during inference. As shown in Table 2, the final evaluation scores are lowest for blood vessels (BLG) at 62.79 %, followed by capillaries (KAP) at 75.62 % and astrocytes (AST) at 89.22 %. For the remaining classes, the classifier achieves high values above 90 %.

**Table 2.**
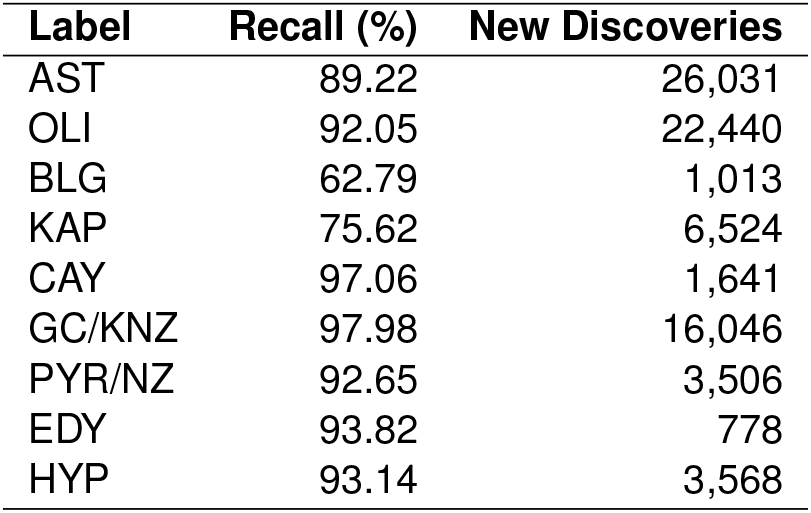
Validation set results: Recall analysis and identification of new discoveries per cell class.

**Table 3.** Detailed definitions and histomorphological features of the annotated and trained cell types and auxiliary technical classes.

| ID | Label | Cell Class | Description |
| --- | --- | --- | --- |
| 1 | BGR | Background | Technical auxiliary class representing the neuropil and extracellular matrix, appearing as a pale violet to bluish ground substance in hematoxylin staining. |
| 2 | AST | Astrocyte | Star-shaped glial cell responsible for maintaining the blood-brain barrier, providing neuronal metabolic support, and mediating reactive gliosis following injury. |
| 3 | OLI | Oligodendrocyte | Specialized glial cell type that produces myelin sheaths. |
| 4 | BLG | Blood Vessel | Larger tubular structures (arteries or veins), responsible for regional blood transport. |
| 5 | KAP | Capillary | Microvascular vessels consisting of a single endothelial layer. |
| 6 | CAY | Corpora Amylacea | Small, spherical glycoprotein aggregates that accumulate during aging. |
| 7 | GC | Granule Cell | Small, densely packed neurons primarily located in the dentate gyrus of the hippocampus. |
| 8 | PYR | Pyramidal Cell | Large, triangular projection neurons acting as the primary excitatory cells. |
| 9 | EDY | Ependymal Cell | Ciliated epithelial cells forming a monolayer that lines the ventricular system. |
| 10 | HYP | Hypoxic Cell | Neurons exhibiting perisurgical damage due to oxygen deprivation. |
| 11 | DN | Dysmorphic Neuron | Pathologically altered neuron featuring an enlarged cell body and aberrant dendritic branching, characteristic of focal cortical dysplasia ILAE Type II. |
| 12 | BC | Balloon Cell | Large, rounded cell with opalescent cytoplasm, serving as a diagnostic hallmark of FCD II. |
| 13 | UNKW | Unknown | Technical auxiliary class utilized to enhance the fundamental object detection and generalization capabilities of the computational model. |

### Statistics

We use the two-sided Mann-Whitney-Wilcoxon test for our comparison tests. We chose this nonparametric test because not all of the variables being tested were normally distributed. The following figures denote statistical differences: ns for p *>* 0.05, * for p ≤ 0.05, ** for p ≤ 0.01, *** for p ≤ 0.001 and **** for p ≤ 0.0001. We conduct comparison tests using the statsannotations Python library [17]. We conduct correlation analysis using Kendall’s *τ* , for which we use the implementation of the pandas Python library [71].

## Results

### A two-term Ogden model well captures the mechanical response of surgical brain tissue

To quantify the mechanical properties, we fit the modified two-term Ogden model defined in Equation 3 to the measured and postprocessed mechanical response for each specimen. We use the approximated hyperelastic response of the third compression-tension cycle and the third torsional shear cycle in the first load step up to a shear of 0.15, shown in Figure 2 c,d. Figure 5 shows the models capability to well capture the experimental data for an exemplary specimen. The compression-tension data in Figure 5 a visualizes the pronounced compression-tension asymmetry with a ratio of maximum absolute stress values 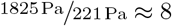 due to the strong strain stiffening in compression, characteristic for brain tissue [14]. This behavior is also reflected by the obtained model parameters with large negative values for *α*_1_. The comparably small strain stiffening in tension is indicated by positive values *α*_2_ *>* 0. Lacking strain stiffening in compression or tension would be indicated by *α*_1_ = 0 or *α*_2_ = 0,respectively, as shown in Figure 3. The response of a linear model shown in Figure 5 a starts to deviate from the experimental data for compressive loading below stretch values of *λ* ≤ 0.95. For the second loading mode, torsional shear shown in Figure 5 b, the model is also able to capture the experimental response with the same set of parameters. This demonstrates the models ability to capture the mechanical behavior under varying loading modes.

**Figure 5.**
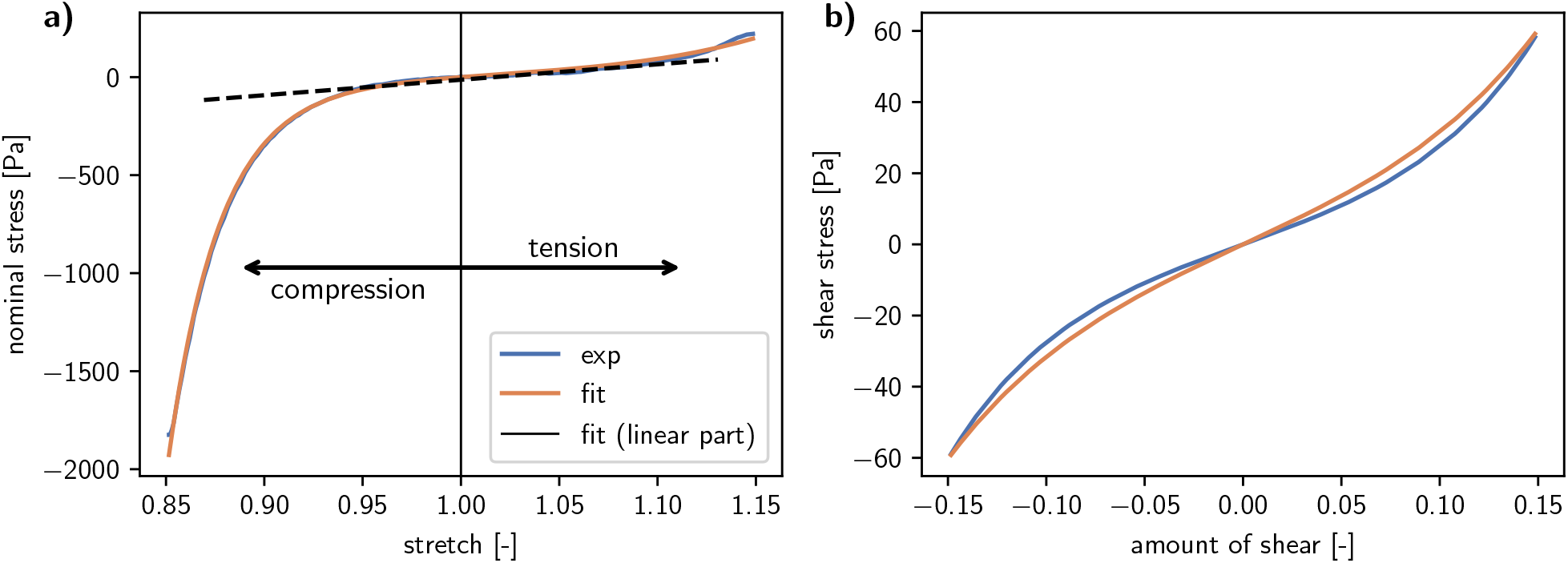
Measured mechanical response (exp; blue) that is well captured by the predicted model output (fit; orange) with identified parameters *α*_1_ = −31.36, *α*_2_ = 18.72 and *µ* = 258.45: a) compression-tension loading from the third cycle with compression-tension asymmetry due to pronounced strain-stiffening in compression. The dotted line represents the predicted response of a linear elastic model with the same shear modulus as the fitted two-term Ogden model. b) Symmetric torsional shear loading in positive and negative direction

### Hippocampal sclerosis specimens show distinct nonlinear stiffening under compression

In this work, we aim to identify characteristic mechanical differences in hippocampal sclerosis tissue. To this end, we have tested 7 specimens from surgically resected tissue of confirmed HS patients as well as 21 specimens extracted from brains of body donors. We have then obtained mechanical parameters *α*_1_ and *α*_2_, characterizing the strain stiffening in compression and in tension, respectively, and the shear modulus *µ*, characterizing the linear contribution of the mechanical response. For small deformations, the nonlinearity introduced by the *α* parameters can be neglected and the response described by *µ* considered sufficient. Parameters were obtained for each specimen. The resulting dataset of parameters in Figure 6 enables us then the systematic comparison of hippocampal specimens with and without confirmed HS diagnosis. We observe consistently lower *α*_1_ values for HS specimens in Figure 6 a which are confirmed as significant by pairwise tests. This means, that HS specimens behave stiffer under high compressive deformations within our tested strain amplitude up to 15 %. For *α*_2_, both groups contain values of *α*_2_ = 0 in Figure 6 b, indicating the tension stiffening contribution being inactive. Mean values of *α*_2_ are higher for body donor specimen but the overall range of obtained value largely overlaps and no significant difference is found. Mean values for the shear modulus *µ* in Figure 6 c are higher for HS specimen but again the value ranges largely overlap and no significant difference is found. In summary, the analysis of mechanical properties reveals strong separation for large compressive deformation due to the compression stiffening captured by *α*_1_ while *α*_2_ and *µ* lie in similar ranges with no significant difference.

**Figure 6.**
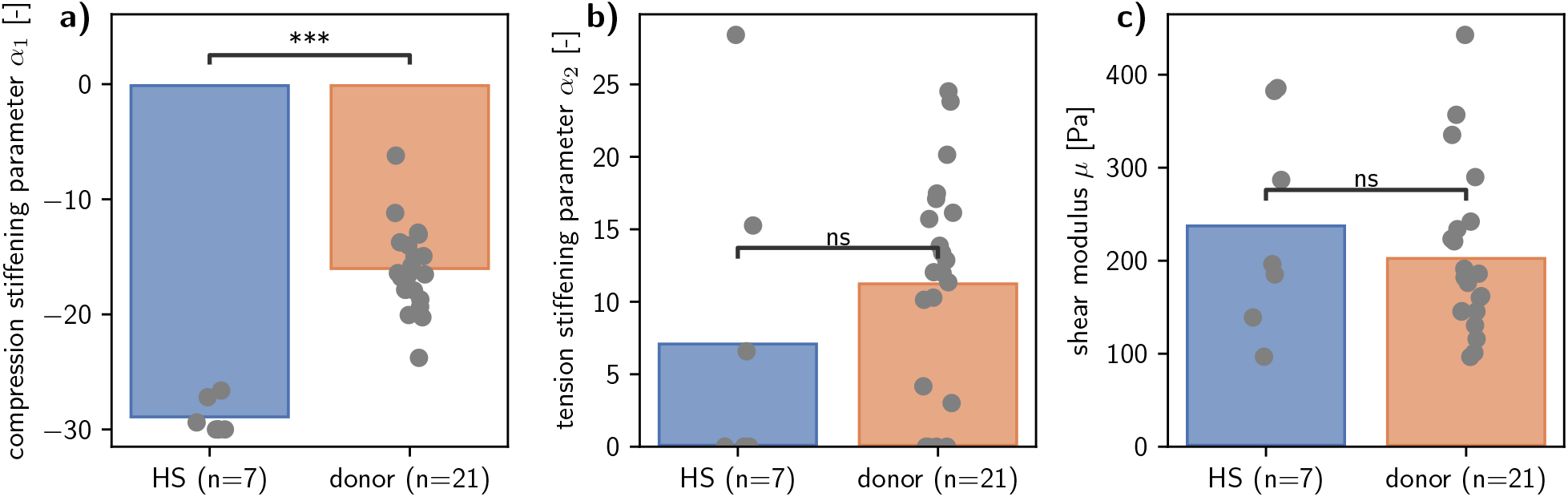
Model parameters for Equation 3 obtained by fitting the measured data of each specimen of surgical resected hippocampal sclerosis tissue (HS,blue) and hippocampus tissue of human body donors (donor,orange). Bars show mean values and black dots denote single specimens: a) *α*_1_ values characterizing the strain stiffening under compression. b) *α*_2_ values characterizing the strain stiffening under tension. c) The shear modulus *µ* characterizing the linear part of the mechanical response for small deformations.

### Nerve cell depletion in hippocampal sclerosis is detected

Our deep neuopathology classifier enables us the automated detection and classification of cells in hematoxylin WSIs. This enables us to analyze the cellular microstructure of our tested specimens and investigate if neuropathological hallmarks of HS can be recognized from the classification results. By considering all detected cells, irrespective of their detected type, we can calculate the overall cell density using the tissue area contained in the WSIs. Comparing the overall cell density for HS and body donor specimens in Figure 7a shows higher mean values for HS specimens which are confirmed as statistical significant by a pairwise test. These differences persists when we consider only glia cells and their density in Figure 7 b, which is again confirmed as statistical significant. No significant difference is found for the neuron density in Figure 7 c. Instead of looking at different cell types independently we can also infer important information by combining them. Here, we use the ratio of detected neuron to glial cells that we choose as particularly suitable to characterize the death of nerve cells characteristic for HS. Consistent with this, the resulting nerve to glia ratio values in Figure 7 d show lower values for HS specimens which are also confirmed as statistically significant.

**Figure 7.**
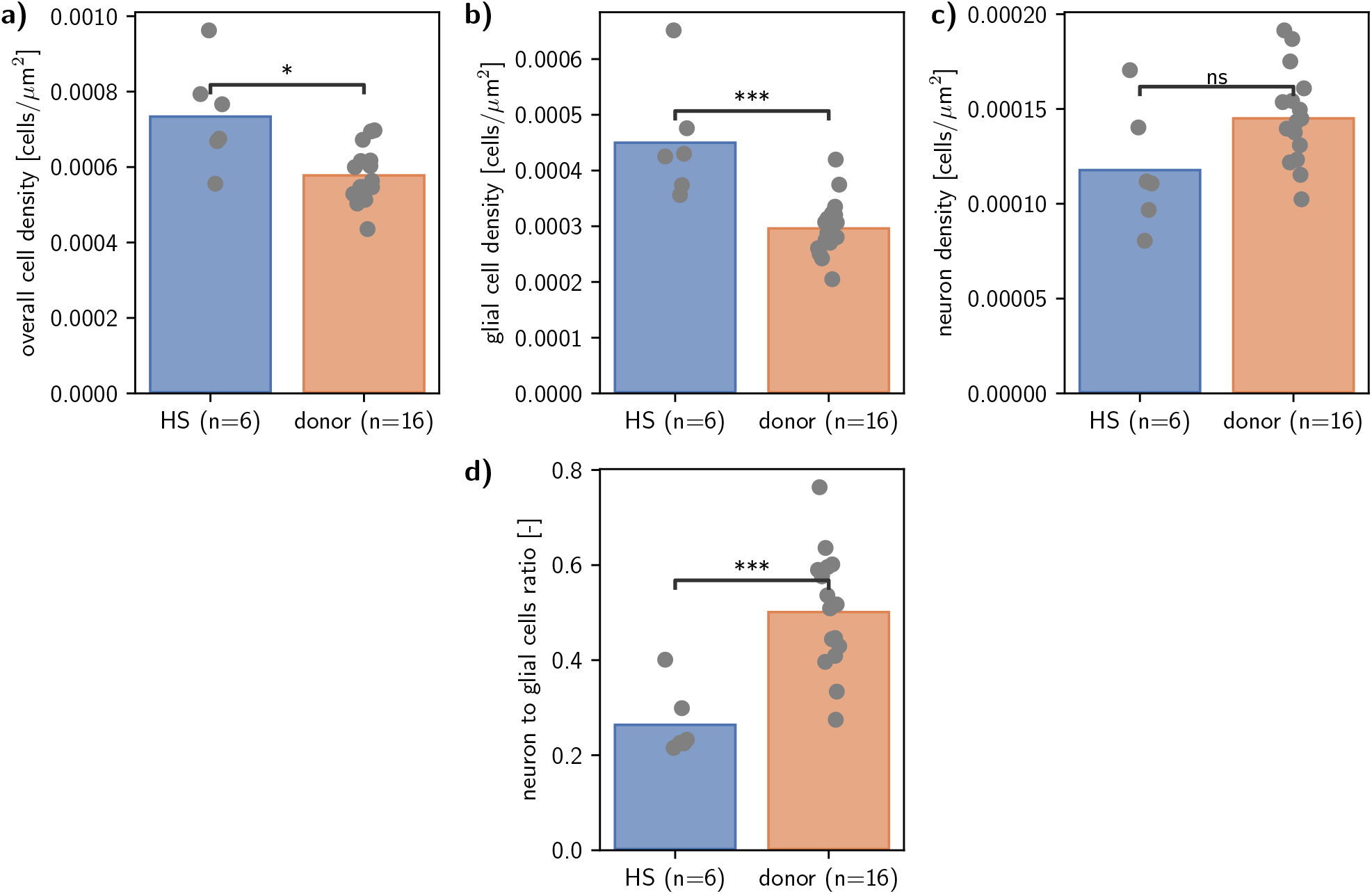
Cellular density in hippocampal sclerosis (HS, blue) and body donor (donor, orange) specimens. Black dots denote single specimens.: a) Overall cell density summing up all cell detections. b) Density of glia cells. c) Density of neurons. d) Ratio of number of detected nerve to glia cells for each specimen.

### Cellular density correlates with tissue stiffness

So far, we have analyzed the mechanical properties and cellular densities independently of each other. By combining them, we can identify general trends that link changes in the observed mechanical response to changes in the tissue microstructure. The scatter plots in Figure 8 show data points for the shear modulus *µ* over different cellular microstructure quantities, as we observe strong relations only for the shear modulus *µ*. The displayed linear regressions capture general trends by taking into account all data points from HS and body donor specimens, where we quantify the identified trends with the annotated Kendall’s rank correlation coefficient *τ* . Figure 8 a shows that for specimens with higher overall cell density we identify higher shear moduli *µ* indicated by a positive correlation coefficient *τ* = 0.44. When we consider the density of only glial cells we identify the same trend in Figure 8 b with a similar correlation coefficient *τ* = 0.39. For the neuron density in Figure 8 c no clear trend is identified as indicated by a low correlation coefficient *τ* = 0.04. Figure 8 d well aligns with the previous observations as the low ratio of neuron to glial cells, indicating nerve cell depletion, together with higher shear moduli *µ* in HS specimens lead to an overall negative trend where we observe lower shear moduli *µ* for higher ratios of neuron to glial cells. Still, this trend is less pronounced, indicated by lower absolute values for the calculated correlation coefficient *τ* = −0.26. The combination of the compression stiffening, quantified by *α*_1_, and the neuron to glial cells ratio in Figure 9 shows two distinct clusters for HS and donor specimens.

**Figure 8.**
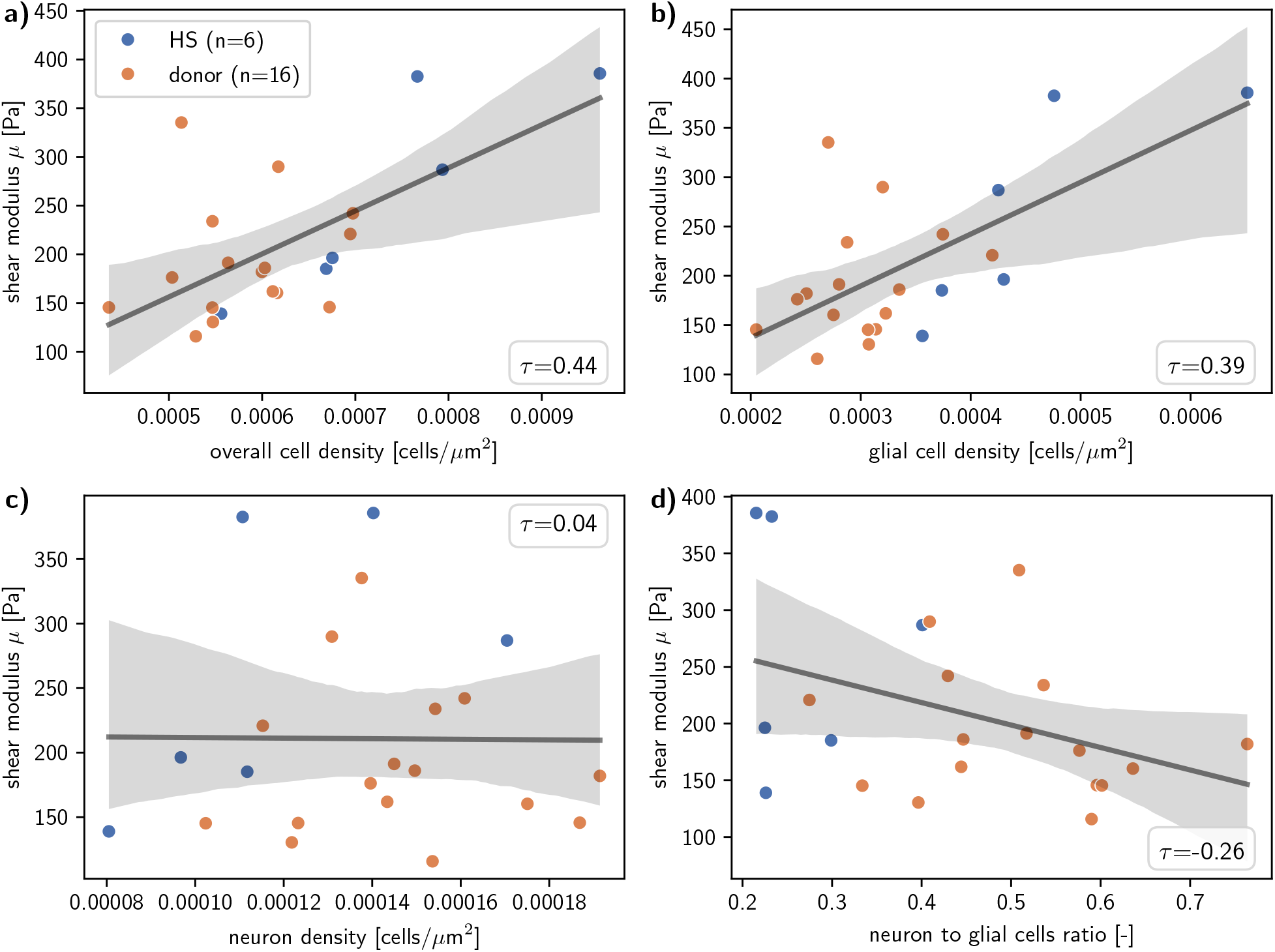
Cellular densities in relation to mechanical parameters. Data points show hippocampal sclerosis (HS, blue) and body donor (donor, orange) specimens. The black line shows the linear regression over all data points. Grey shadows show the 95% confidence interval. *τ* values indicate Kendall’s rank correlation coefficient *τ* .

**Figure 9.**
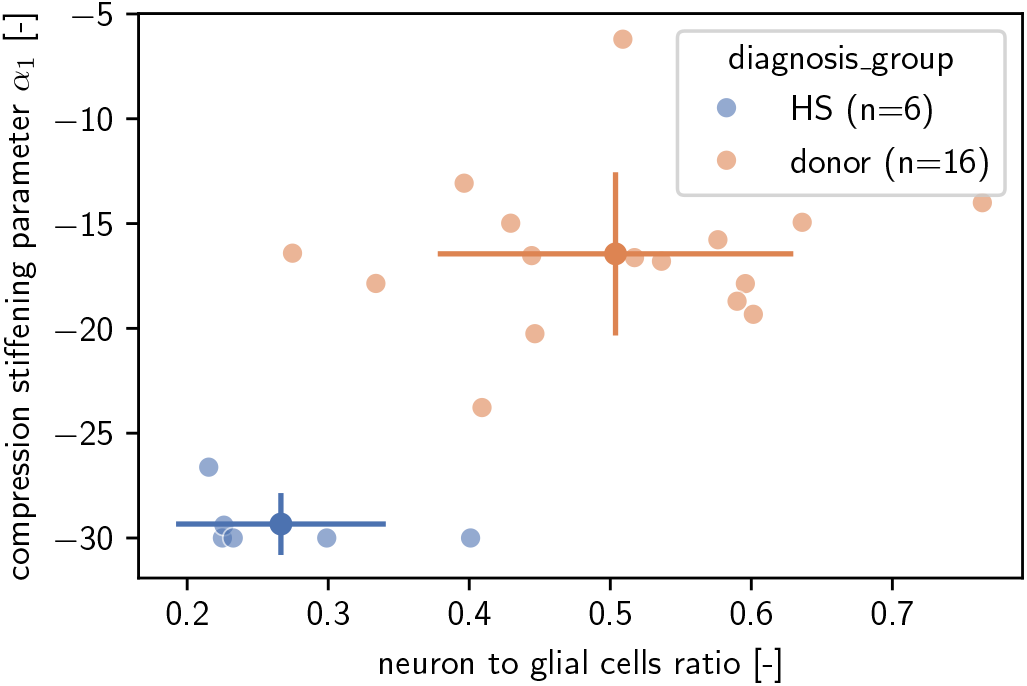
Mechanical parameters *α*_1_ characterizing strain stiffening in compression over the ratio of detected nerve to glia cells for hippocamapal sclerosis specimens (HS, blue) and body donor specimens (donor, orange). Bold dots denote mean values with lines marking the standard deviation and light colored dots single measurements.

### Hippocampi volume ratios align with detected nerve cell depletion

An important method in the diagnosis of HS is magnetic resonance imaging (MRI). One of the characteristic hallmarks that can be inferred from MRI data is the reduction of the hippocampus volume in HS [49]. Here, we have extracted hippocampi volumes from MRI data available for the HS patients where we also tested surgically resected tissue samples. All patients had only one hemisphere affected by HS, enabling us to derive the volume ratio of unaffected to affected hippocampus volume as metric for disease progression. The positive relation of neuron to glial cells ratio to the volume ratio in Figure 10 is confirmed by Kendalls correlation coefficient with a value of *τ* = 0.6. This result validates the ratio of neuron to glial cells, obtained from our deep neuropathology classifier and measuring nerve cell depletion in HS, as marker for HS progression.

**Figure 10.**
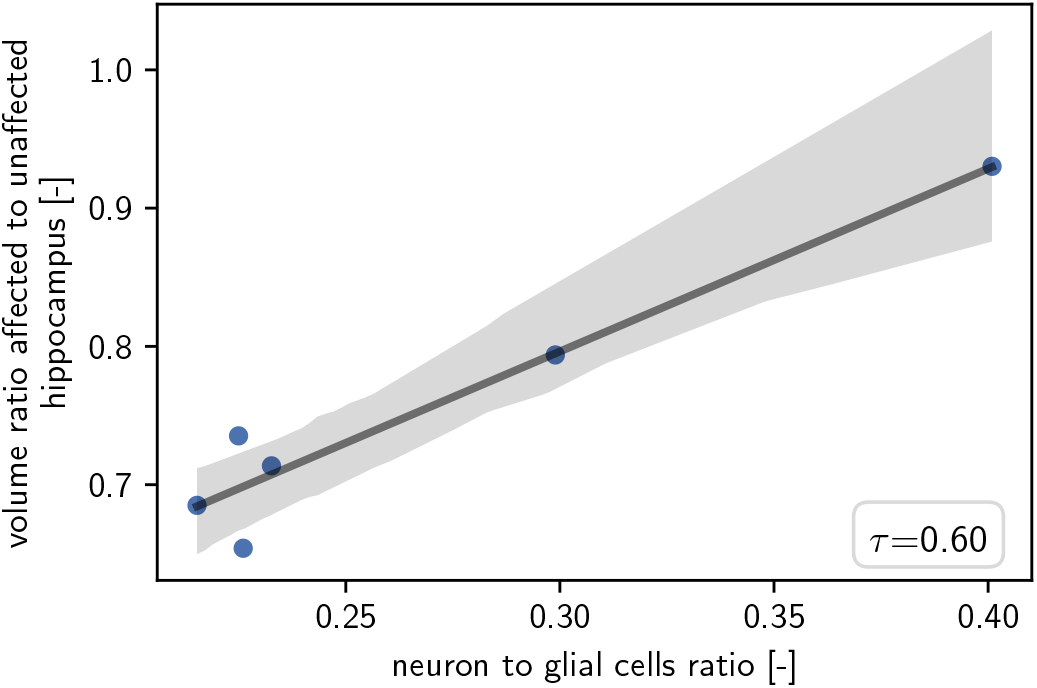
Hippocampus volume ratio of affected to unaffected hemisphere over ratio of detected neuron to glial cells. Only data points from HS specimens are shown. The black line shows the linear regression over all data points. Grey shadows show the 95% confidence interval. *τ* value indicates Kendall’s rank correlation coefficient

### Fractional anisotropy as an *in vivo* measurement for stiffness

Besides the hippocampus volume utilized in the previous section, the fractional anisotropy (FA) is also obtained from MRI data and commonly used in HS diagnosis. FA values are calculated from diffusion tensor imaging (DTI) sequences and indicate the isotropy of the water diffusion. They range from 0 -purely isotropic- to 1 -fully anisotropic with diffusion constrained to one direction- and have shown negative correlation with tissue stiffness in previous studies [65, 14]. Here, we relate the shear modulus *µ* and the FA value averaged over the hippocampus. Importantly, the data points here only partially overlap with the patients and donors of the previous shown mechanical and cell density data as FA measurements were only available for a subset of HS patients (n=2) and body donors (n=10). Considering all data points in Figure 11 we do not find a clear relation of FA to *µ* values. The neuropathological analysis of the tested donor brains revealed substantial Alzheimer’s disease related findings in two brains. Therefore, we exclude these data points as they are not representative for this study, focusing on HS specific differences, and find a strong negative correlation indicated by Kendall’s correlation coefficient *τ* = − 0.62. Thereby, we show that the relation of FA measurements and mechanical behavior -in terms of the small deformation response characterized by the shear modulus *µ*-extends beyond previously identified overall trends for region-specific parameters average over multiple brains to inter-individual trends for the same region – the hippocampus.

**Figure 11.**
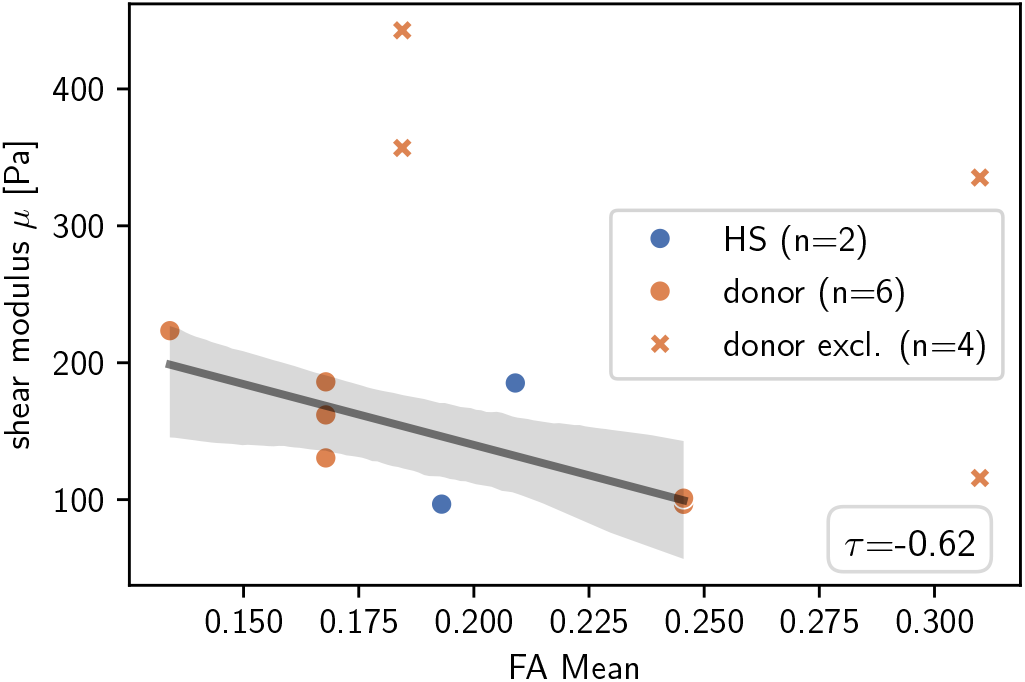
Shear modulus *µ* over the averaged fractional anisotropy (FA). Data points show hippocampal sclerosis (HS, blue) and body donor (donor, orange). X marker denote specimens not considered for the linear regression. The black line shows the linear regression over all data points. Grey shadows show the 95% confidence interval. *τ* values indicate Kendall’s correlation coefficient.

## Discussion

In this study, we have characterized the mechanical behavior of the sclerotic hippocampus by mechanically testing surgically resected tissue with histopathologically confirmed HS diagnosis. As a control group, we additionally tested tissue from the hippocampi of body donors. For each specimen, we quantified nonlinear material parameters using a hyperelastic, two-term Ogden model. While characterizing the mechanical properties enables us to identify characteristic behaviors of pathological brain tissue, it does not provide any insight into underlying microstructural changes. Therefore, we have performed histopathological and diffusion-weighted MRI analyses to additionally quantify the underlying cellular microstructure and composition. Combined analysis of mechanical properties and cellular composition allows us to identify general relations between the two domains and to clearly distinguish between the healthy and pathological conditions. This is an important step towards mechanics-based diagnostic biomarkers.

### Hippocampal sclerosis tissue shows significant increase in compression stiffening under large deformations

In this study, we characterized the mechanical behavior of human brain tissue from the hippocampus with and without hippocampal sclerosis. To this end, we combined large strain *ex viv* o mechanical testing under compression, tension, and torsional shear with nonlinear continuum mechanics modeling. The mechanical response initially follows an approximately linear stress-strain relationship, indicated as dotted line in Figure 5. However, clear deviations from the linear approximation are observed when the sample is further deformed beyond 5% strain. This highlights the importance of accounting for nonlinear effects when the goal is to quantify mechanical properties relevant for neurosurgical intervention, where large deformations occur. To this end, we fitted a modified two-term Ogden model to the measured response and obtained three parameters for each specimen: the shear modulus *µ* and the strain stiffening parameters *α*_1_ and *α*_2_. For the linear contribution, characterized by *µ* and visualized in Figure 5, no significant differences were found between the HS and body donor groups in Figure 6. Nevertheless, we identified a clear mechanical distinction between the two groups in the form of consistently lower *α*_1_ values, which characterize strain stiffening under compression. These differences were confirmed as statistically significant.

In general, quantitative mechanical properties of epileptic brain tissue are still sparse in the current literature. Gautam et al. [25] conducted *ex vivo* nanoindentation on freshly resected tissue with and without epilepsy and reported that the tissue with epilepsy exhibited a stiffer response, as indicated by higher storage moduli (G’). Within the epilepsy group, they obtained the stiffest values for tissue from patients with mesial temporal sclerosis. While these results are not confirmed by our data, which show no significant difference in shear modulus between healthy subjects (HS) and donor specimens, the comparison is somewhat limited as the extraction location of the brain tissue is not specified. Another study using *ex vivo* indentation tested only tissue from patients with mesial temporal lobe epilepsy and obtained shear modulus values within the range reported for healthy brain tissue [59].

*Ex vivo* AFM tests of dentate gyrus granule cells in the hippocampi of an epilepsy mouse model showed increased stiffness compared to wild type [76]. Our findings do not confirm these results, although a direct comparison is limited by differences in scale (tissue vs. cell level) and species (human vs. mouse).

Huesmann et al. [36] obtained *in vivo* measurements of the hippocampus tissue of epilepsy patients via magnetic resonance elastography (MRE). Intraindividual comparisons revealed higher shear modulus *µ* values for the ipsilateral hippocampus compared to the contralateral one, with a statistically significant overall difference in the intraindividual stiffness ratio. The shear modulus values *µ* obtained in the present study do not show this distinct difference. This could be due to differences in the boundary conditions between *in vivo* MRE and our *ex vivo* rheometer measurements. Tissue perfusion, vasculature, support, and interstitial fluid flow due to surrounding tissue may contribute to these differences. Shear wave elastography (SWE), closely related to MRE, is suitable for intraoperative mechanical measurements. It has been successfully applied in epilepsy surgery to identify focal lesions, as it showed distinct stiffness values between healthy brain tissue and lesion sites [51, 16]. However, SWE data for HS tissue are not available in the literature, preventing direct comparison to our results.

That being said, MRE, SWE, and *ex vivo* finite strain measurements could serve as complementary methods for measuring the mechanical properties of hippocampal tissue under different conditions, contributing to a complete mechanical characterization of brain tissue.

### Gliosis relates to tissue stiffening in hippocampal sclerosis

Our observation that sclerotic hippocampus tissue exhibited a significant increase in compression stiffness raises the question of how mechanical changes at the tissue level are linked to underlying neuropathological changes at the cellular level. To investigate this, we have implemented a deep neuropathology classifier that uses a neural network to detect and classify neuronal and glial cells in whole slide images (WSI) of hematoxylin-stained tissue slices. We applied the classifier to WSIs of the mechanically tested tissue and obtained cellular densities, providing detailed information on cellular composition. This allowed us to identify the relationship between cellular composition and the linear contribution to the mechanical response, as measured by the shear modulus *µ*. Figure 8 a shows a positive correlation between *µ* and overall cell density.

Using our cell classifier, we were able to distinguish different cell types and obtain neuron and glial cell densities individually. We observed the same trends for glial cell density as for overall cell density in Figure 8 b, where samples with higher cell densities also exhibited higher shear moduli *µ*. However, we did not find a correlation between neuron density and shear modulus *µ* in Figure 8 c. Still, by combining glial cell and neuron detection counts to calculate a neuron-to-glia ratio, we observed a weak correlation with *µ* in Figure 8 d that also links linear mechanical properties to an HS marker.

Cell densities are obtained as the ratio of detected cells to the tissue area of the whole slide image (WSI). Tissue area relates to volume, and volume loss (atrophy) is a well-known characteristic of hippocampal sclerosis. For a sclerotic hippocampus, a correlation between decreased volume and neuronal cell count has been demonstrated [10]. This volumetric loss could explain the lack of significant differences in neuronal density between HS and donor specimens in our cell density results (see Figure 7). In this case, the loss of neurons might be countered by the loss of volume. Accordingly, this would also explain why we did not observe a correlation between shear modulus and neuron density.

The positive correlation between cellular density and tissue stiffness is not universal to CNS tissue, as studies investigating the brain tissue mechanics of different species have shown. A positive relationship between stiffness measurements and overall cell density was found using atomic force microscopy (AFM) in developing Xenopus laevis [4], mouse spinal cord [43], and indentation of mouse hippocampus and cerebellum [2]. On the contrary, our previous findings from *ex vivo* experiments on human brain tissue, in which we compared different regions (basal ganglia, corpus callosum, corona radiata, and cortex), show higher shear moduli in regions with lower overall cell density [11]. In another study by our group on porcine brain tissue, we found the same relationship for the equilibrium response parameters representing the hyperelastic contribution of a viscoelastic model [67]. However, both studies investigated interregional trends (excluding the hippocampus) without looking at pathological changes, whereas here, we tested hippocampal tissue with and without HS.

In addition to neuropathological analyses, we used pre-resection diffusion tensor imaging (DTI) to obtain fractional anisotropy (FA) values, indicating the directionality of nerve fiber bundles in the tissue, for each mechanically tested hippocampus. Figure 11 shows a negative correlation between FA values and the shear modulus *µ*, which aligns well with previous studies [65, 14].

The characteristic stiffening of the sclerotic hippocampus contrasts with the softening observed in acute central nervous system (CNS) injuries, as observed in rats [55] and an Alzheimer’s disease model in mice [27]. All of these pathologies are accompanied by gliosis, and single-cell atomic force microscopy (AFM) measurements support gliosis as a softening mechanism, as glial cells were found to be consistently softer than neurons [47]. These seemingly contradictory results could be explained by differences in observation time points. Studies on the rat spinal cord 12 weeks post-injury measured by AFM [21] and indentation [37] report stiffer elastic moduli compared to healthy controls. Cooper et al. found fibrotic scar components to be a likely explanation for the stiffening [21]. We hypothesize that changes in the extracellular matrix (ECM) driven by gliosis contribute to stiffening during chronic hippocampal sclerosis pathology. Nevertheless, further investigation of ECM changes and their relation to mechanics in HS pathology is needed.

### Mechanics augmented surgery: towards *in vivo* diagnostic tools

In order to use mechanical cues for pre- and intraoperative planning, it is necessary to translate mechanical measurements to the clinical context. One available approach is magnetic resonance elastography (MRE), which is suitable for preoperative planning and has been shown to identify for example tumors [15]. Another elastography method is shear wave elastography (SWE), which is suitable for intraoperative applications and has been used for e.g. epilepsy lesion localization [52]. However, elastography methods are currently limited to small deformations and cannot reach the loading level at which the stiffening effects observed in this study distinguish sclerotic from healthy hippocampus. Additionally, difficulties obtaining sufficient signal strength from deeper brain regions, such as the hippocampus, can pose additional challenges for elastography methods. “Smart” instruments like ultrasonic aspirators enable targeted measurements but are limited to small strains [8].

One promising approach to integrate mechanical cues into surgical procedures is the use of tactile surgical tools that measure the forces exerted by the surgeon on the tissue [63, 64, 29]. These forces, when combined with the resulting tissue deformation, enable the estimation of *in vivo* tissue stiffness, which can then be compared with pathological thresholds. Further research is necessary to confirm our *ex vivo* findings for the sclerotic hippocampus in an intraoperative setting and to determine such mechanical thresholds.

### Limitations

We have tested small cylinders of brain tissue *ex vivo* by submerging and gluing them to the specimen holders of a rheometer. This setup allows us to measure the tissue’s mechanical response under different loading conditions and determine its complete mechanical profile. However, differences remain in terms of the missing boundary conditions of surrounding tissue, vasculature, blood pressure, and interstitial fluid flow compared to *in vivo* or *in situ* conditions. Therefore, the combination of different mechanical measurement approaches is crucial to obtaining a complete picture of brain mechanics.

Further limitations arise from differences between HS and body donor groups beyond the existence or lack of an HS pathology. Surgical tissue was tested within approx. six hours after resection while in the case of body donor brains, time required for transport, brain extraction, and MRI scanning led to post mortem times of up to 89 hours when the tissue was tested. Another factor is the age difference between the two groups, with patients ranging from 21 to 55 years and body donors ranging from 57 to 92 years. Especially in donors on the upper end of the age range, we found neuronal cell loss and plaques in the tested HS samples, which may have influenced our results.

## Conclusion

In this study, we identified distinct differences in the nonlinear mechanical behavior of the hippocampus in cases with and without hippocampal sclerosis (HS). Surgically resected HS tissue showed higher stiffness under *ex vivo* compressive loading compared to hippocampus from body donors. Thus, we quantified the characteristic firmness of HS tissue as perceived by neurosurgeons and neuropathologists. The obtained mechanical parameters are an important contribution to translating mechanical cues into intrasurgical diagnostic approaches for epilepsy surgeries. To establish a connection between mechanics and cellular composition, we implemented a cell detection and classification pipeline that utilizes a neural network to automatically process WSIs. In HS samples, we observed a significant increase in glial cell density and a significant reduction in the neuron-to-glial cell ratio. This indicates gliosis and neuronal cell loss in HS surgical specimens, respectively. These neuropathological hallmarks align with the histopathological criteria for a HS diagnosis. Thus, the classifier results confirm the histopathological diagnosis. Additionally, we validated our cell detection results using MRI-derived hippocampal volume ratios, which indicate HS-induced atrophy and are clearly related to the neuron-to-glial cell ratio.

We further identified relationships between the linear mechanical response, as defined by the shear modulus *µ*, and cellular composition. Specimens with higher overall density exhibited stiffer responses. The same trend was found for glial cell density, which supports the hypothesis of a gliosis-driven tissue stiffening mechanism.

## AUTHOR CONTRIBUTIONS

S.B. and I.B. conceptualized the study and acquired funding. S.B. and I.B. supervised the study. S.B., I.B. J.V., N.R., L.H., S.R. and J.H. wrote the initial manuscript draft. N.R. and J.H. conducted mechanical experiments. J.H. implemented the code for the mechanical model, analyzed the data and visualized the results. I.B. and L.H. conducted neuropathologocial examinations of tissue specimens. I.B. annotated cell classes for the training of the classifier. D.D. and O.S. provided surgically resected tissue. J.V. implemented the cell classifier, evaluated whole slide images, analyzed the data and visualized results. S.R. conducted MRI measurements for epilepsy patients and body donors, and analyzed and processed the data. L.B. and F.P. provided human body donors. All authors discussed the results, reviewed, and edited the manuscript.

## ACKNOWLEDGEMENTS

We gratefully acknowledge the financial support by the Deutsche Forschungsgemeinschaft (DFG, German Research Foundation) through project 460333672 CRC1540 EBM (project A01 and A02). The authors wish to sincerely thank those who donated their bodies to science so that anatomical and biomechanical research could be performed. Results from such research can potentially improve patient care and increase mankind’s overall knowledge. Therefore, these donors and their families deserve our highest gratitude. We would like to thank Lisa Stache for preparing human brains from body donors.

## COMPETING FINANCIAL INTERESTS

The authors declare no competing interests.

## DATA AVAILABILITY STATEMENT

The code generated and analyzed during the current study as well as the experimental data is available from the corresponding author on reasonable request.

## Supplementary Information

### A. Cell classifier

1 http://neuroimage.usc.edu/brainstorm

## Notes

### Competing Interest Statement

The authors have declared no competing interest.

